# Natural variation in *Arabidopsis* ISR1 affects iron localization and induced systemic resistance

**DOI:** 10.1101/2021.09.01.458588

**Authors:** Amanda L. Socha, Yi Song, Brandon S. Ross, Jenifer Bush, Frederick M. Ausubel, Mary Lou Guerinot, Cara H. Haney

## Abstract

Beneficial root-associated bacteria can induce systemic resistance (ISR) to foliar pathogens and there is known transcriptional and genetic overlap in the root response to iron deficiency and ISR. A previous study found that there is natural variation in ISR among *Arabidopsis* accessions. The Ws accession is deficient in ISR, and the responsible recessive genetic locus, named ISR1, was mapped to chromosome 3. To find candidate genes that may underlie ISR deficiency in Ws, we identified genes that are induced in response to the ISR-triggering bacterium *Pseudomonas simiae* WCS417 and to iron stress and that have non-synonymous mutations in the Ws genome with respect to the ISR-responsive Col-0. We identified a kelch-domain containing protein encoded by *At3g07720* that has a genomic rearrangement in Ws. We found that overexpression of Col-0 *At3g07720* restores ISR to Ws, indicating that *At3g07720* encodes *ISR1. Isr1* loss of function mutants do not affect plant growth under iron limiting conditions but have increased levels of apoplastic iron. We found that iron supplementation, *P. simiae* WCS417, or a loss of *isr1* enhance ROS production in a non-additive manner, suggesting they work through the same mechanism to enhance resistance. Our findings show that *ISR1* is required for iron localization, immunity, and ISR, and suggest that increased iron uptake induced by ISR-eliciting bacteria may directly contribute to immunity through increased reactive oxygen production.

## Introduction

Beneficial root-associated microbes provide diverse benefits to plants ranging from growth promotion to pathogen protection (Berendsen et al., 2012). However, due to the difficulty of carrying out forward genetic screens in many plant-microbiome interaction systems, the genetic and molecular mechanisms by which the host senses and responds to microbiota have remained elusive (Song et al., 2021). Induced systemic resistance (ISR) is a process by which specific members of the root-associated microbiome can enhance above-ground resistance to pathogens. ISR is genetically controlled by the plant host and requires genes that are distinct from those involved in basal immunity in leaves (Pieterse et al., 2014). The genetic basis of ISR in *Arabidopsis* has been most extensively studied using *Pseudomonas simiae* strain WCS417 (Pieterse et al., 2014), but relative to studies on the local response to pathogen infection, the molecular mechanisms that underlie ISR remain relatively unexplored.

There is genetic and functional overlap between the iron deficiency response and ISR by below-ground beneficial microbes. Comparing the transcriptional response of roots to WCS417 on the one hand, and iron deficiency on the other, identified a significant overlap in genes that were upregulated by both *P. simiae* WCS417 and iron limitation (Dinneny et al., 2008; Zamioudis et al., 2015). Although root colonization with ISR bacteria leads to a local iron deficiency response, it can rescue iron deficiency in the shoots (Zamioudis et al., 2015; Trapet et al., 2016). The iron deficiency inducible transcription factor *MYB72* (Palmer et al., 2013) and its target, *BGLU42*, are required for the elicitation of ISR by *Pseudomonas simiae* WCS417 against the foliar bacterial pathogen *P. syringae* (Van der Ent et al., 2008; Segarra et al., 2009; Zamioudis et al., 2014). Treatment by ISR bacteria also triggers secretion of coumarins into the rhizosphere, which can modulate the rhizosphere microbiome (Stringlis et al., 2018; Harbort et al., 2020) and promote iron uptake (Robe et al., 2021). These data indicate that ISR-triggering bacteria modulate both plant iron status and immunity.

Iron is an essential metal micronutrient for eukaryotes as well as their associated pathogenic and beneficial microbes. However, iron levels must be regulated due to the high redox activity of iron, which can lead to the spontaneous formation of reactive oxygen species (ROS) *via* the Fenton reaction (Halliwell and Gutteridge, 1992). The potential of iron to contribute to ROS production as well as its essentiality for life has led to two main theories implicating iron in immunity: 1) the host sequesters iron away from the pathogen to limit microbial growth (Deák et al., 1999; Dellagi et al., 2005; Kieu et al., 2012), and 2) the host utilizes iron to enhance the formation of antimicrobial ROS (Liu et al., 2007; Ye et al., 2014). In plant-pathogen interactions there is evidence for both actively limiting iron availability to restrict pathogen growth, as well as direct utilization of iron to generate ROS (Verbon et al., 2017). Iron accumulation and ROS are observed at the sites of fungal infection by *Blumeria graminis* and *Colletotrichum graminicola* (Liu et al., 2007; Ye et al., 2014) and iron-deficient maize and *Arabidopsis* are significantly more susceptible to pathogen infection (Aznar et al., 2014; Ye et al., 2014). At the same time, iron chelation is a widely used virulence mechanism by bacterial pathogens (Segond et al., 2009; Kieu et al., 2012; Aznar and Dellagi, 2015; Xing et al., 2021). Collectively these data implicate precise regulation of iron availability as critical to the outcome of plant-microbe interactions, indicating that there is tension between the role of iron in defense and a pathogen’s need for iron to survive. Whether a change in iron status triggered by ISR bacteria contributes to enhanced plant immunity is unknown.

Ton et al. previously reported that *Pseudomonas simiae* WCS417 failed to trigger ISR on the *Arabidopsis* accession Ws (Ton et al., 1999; Ton et al., 2001). The *ISR1* locus was mapped to chromosome 3 and the Ws allele was found to be recessive to the Col-0 allele. To identify novel genetic components that link plant iron status with immunity and ISR, we used a reverse genetics approach based on the transcriptional overlap of genes induced during iron deficiency and colonization by *P. simiae* WCS417 (Dinneny et al., 2008; Zamioudis et al., 2015) and looked for genes with non-synomomous mutations within the ISR1 locus. We identified a gene containing a kelch-domain, *At3g07720/ISR1*, which is highly upregulated during iron deficiency and *P. simiae* WCS417 colonization. We show that overexpression of *At3g07720* restores ISR to Ws suggesting that *At3g07720* encodes ISR1. While loss-of-function *isr1* mutants do not have an iron deficiency phenotype, *isr1* mutants have increased levels of apoplastic iron, enhanced immunity, constitutive ISR, and increased ROS, suggesting that *ISR1* plays a key role in immunity, ISR and iron localization.

## Results

### Natural variation in the *At3g07720/ISR1* gene underlies ISR deficiency in the *Arabidopsis* Ws accession

Previous work identified a recessive locus in the *Arabidopsis* Ws accession that is linked to a loss of ISR by *P. simiae* WCS417 (Ton et al., 1999; Ton et al., 2001). To identify candidates for the *ISR1* gene, we used a reverse genetics approach to identify genes that were 1) within the previously described ISR1 locus occuring between the B4 (2.25 Mbp) and GL1 (10.36 Mbp) markers on chromosome 3 (Ton et al., 1999), which encompasses genes At3g07115 and At3g27920, 2) upregulated in the root in response to iron deficiency, and 3) upregulated in roots in response to ISR-activating bacteria based on published datasets (Dinneny et al., 2008; Zamioudis et al., 2015). We identified 8 genes that meet these criteria as strongly induced (log_2_2-fold or greater) in response to ISR bacteria and iron deficiency within the ISR1 locus (Supplementary Table 1).

To further narrow down candidates for the *ISR1* gene, we sequenced the Ws genome using paired end 150 bp Illumina reads, mapped the reads to the Col-0 reference genome and identified non-synonymous mutations within the 8 genes induced in response to iron deficiency and ISR. We identified a ∼72 bp region, encompassing 24 predicted amino acids, with no mapped reads that fell within the predicted open reading frame of WS *At3g07720*, encoding a predicted galactose oxidase/kelch repeat superfamily protein, suggesting a deletion, insertion, or genomic rearrangement (Supplementary Fig. 1). The only other gene among the 8 candidate genes that had nonsynonymous mutations was *At3g10720* encoding a putative plant invertase/pectin methylesterase inhibitor superfamily protein, which had 3 predicted non-synonymous mutations (Supplementary Table 1). To validate the putative deletion in *At3g07720* predicted by Illumina sequencing in the Ws accession, we designed primers to amplify the promoter region, open reading frame, and 3’ end of the predicted open reading frame of *At3g07720* (Supplementary Fig. 1). All primer pairs were able to generate a band of the correct size from Col-0 genomic DNA. While primers amplified the 5’ and 3’ UTR of *At3g07720* from Ws genomic DNA, all primers pairs spanning the 72 bp region failed to amplify from genomic Ws gDNA (Supplementary Fig. 1). As a result, we hypothesized that the genomic changes in this region are more substantial than a 72 bp deletion. We hypothesize that a genomic rearrangement occurred in the Ws accession at this locus and that *At3g07720* may encode ISR1.

We carried out several experiments to determine whether *At3g07720* encodes ISR1. First, we obtained two independent T-DNA insertion mutants in *At3g07720*, one in the Col-0 genetic background and one in the Ws background (Fig. 1A). Using primers that amplify outside of the putative rearrangement in Ws, we tested expression of *At3g07720* under iron deficiency and observed lower induction of *At3g07720* in Ws than in Col-0 (12-fold in Ws versus 25-fold in Col-0; p=0.0007; Fig. 1B). We did not detect *At3g07720* mRNA in the roots of either T-DNA mutant regardless of growth conditions, indicating that these were indeed loss of function mutants (Fig. 1B).

**Figure 1.**
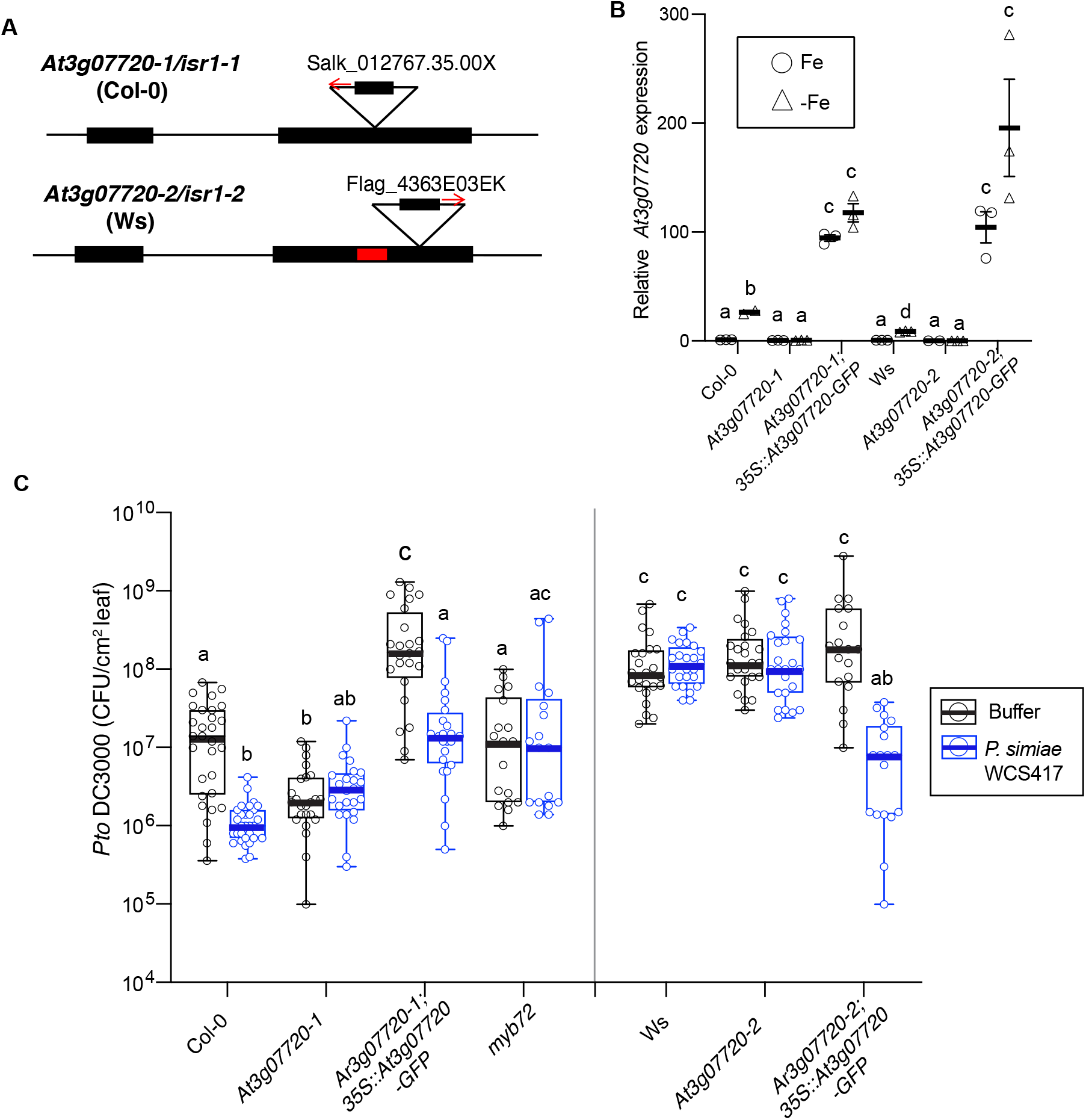
A loss of function of *At3g07720* underlies the ISR-deficient phenotype in Ws. A, Two T-DNA insertion alleles in *At3g07720*/*ISR1* designated *At3g07720-1*/*isr1-1* (Col-0 background) and *At3g07720-2/isr1-2* (Ws background) were used in this study. The red insertion in the Ws allele denotes the location of a putative genomic rearrangement. B, Quantitative RT-PCR Analysis of *ISR1* transcript levels relative to *Clatherin* in roots of plants grown for 10 days and transferred to 50 µM iron (+Fe) or 300 µM ferrozine (-Fe) plates for 3 days. C, Phenotypes of T-DNA lines of insertions in *Arabidopsis ISR1* in Col-0 or Ws backgrounds. While Col-0 shows ISR against *Pto* DC3000 in the presence of *P. simiae* WCS417, the Col-0 *myb72* and *isr1-1* mutants are defective in ISR. Ws was previously shown to be defective in ISR. Overexpression of the *ISR1* gene restores ISR in the Ws ecotype. For all experiments, plants were grown on Jiffy pellets for 5 weeks before infiltrating adult leaves with *Pto* DC3000. CFUs/cm^2^ leaf tissue were quantified 2 days after bacterial challenge. B-C, Letters indicate statistical significance p<0.01 by ANOVA and Tukey’s HSD tests from 3 independent experiments with individual data points shown.

Second, we expressed the Col-0 allele of *At3g07720* tagged with GFP in Ws driven by the 35S promoter. These plants constitutively expressed *At3g07720* regardless of iron status (Fig. 1B). We then tested these transgenic plants as well as the *At3g07720* T-DNA mutant plants described above for their ability to exhibit ISR. Specifically, we treated the roots of soil-grown *Arabidopsis* seedlings with the ISR-inducing bacterial strain *P. simiae* WCS417 and challenged the leaves of adult plants with *Pseudomonas syringae* pv. tomato DC3000 (*Pto* DC3000) (Cecchini et al., 2019). While we observed ISR against *Pto* DC3000 on Col-0 plants treated with *P. simiae* WCS417, we found that the Col-0 *myb72* mutant, both *At3g07720* T-DNA insertion lines, and the Ws ecotype were completely deficient in ISR (Fig. 1C). However, constitutive overexpression of the full-length *At3g07720* gene restored ISR in Ws, indicating that *At3g07720* can complement the ISR-deficient phenotype in Ws (Fig. 1C). Collectively these data indicate that *At3g07720* encodes *ISR1*.

The Ws accession is more susceptible than Col-0 to *Pto* DC3000 infection (Ton et al., 2001). In contrast, we observed that insertions in Col-0 *ISR1* resulted in a 3-4-fold decrease in *Pto* growth compared to wild type Col-0 (Fig. 1C). While we found that the Ws accession was more susceptible than Col-0 to *Pto* DC3000 infection, confirming the results of Ton et al (1999, 2001), an insertion in the Ws *ISR1* gene did not further affect *Pto* DC3000 growth. Conversely, *35S:ISR1-GFP* expressing lines were significantly more susceptible to *Pto* DC3000 in both the Ws and Col-0 genetic background (Fig. 1C). These data suggest that *ISR1* negatively regulates *Arabidopsis* resistance to *Pto* DC3000 but that a loss of *At3g07720* function does not explain the enhanced susceptibility of the Ws accession. This observed role of *ISR1* is similar to the role of jasmonic acid signaling as a positive regulator of ISR and a negative regulator of immunity (van Wees et al., 2000).

The *ISR1* locus was previously shown to be linked to ethylene sensitivity in the Ws ecotype (Ton et al., 2001). However, when we tested the parental Col-0 and Ws lines as well as *isr1-1* (Col-0) and *isr1-2* (Ws) T-DNA insertion mutants, we found no changes in ethylene sensitivity (Supplementary Fig. 2). These data indicate that the ethylene sensitivity phenotype in Ws is distinct from an *isr1* null phenotype and that the ethylene sensitivity phenotype previously reported in ecotype Ws was either lost in the Ws accession used in this study or is due to genetic differences in accessions designated “Ws” across labs (Anastasio et al., 2011).

### Loss of function of *At3g07720/isr1* results in increased levels of apoplastic iron

Our data indicate that *At3g07720* encodes *ISR1* and that natural variation in this gene contributes to natural variation in ISR in *Arabidopsis* (Fig. 1; Supplementary Fig. 1). *At3g07720* has been proposed to be the ancestor to *Arabidopsis* nitrile-specifier proteins (NSPs) (Kuchernig et al., 2012), which are involved in breakdown of glucosinolates in *Arabidopsis* (Kong et al., 2012). NSPs utilize iron as a cofactor, and although *At3g07720* (aka *AtNSP1-like*) has been shown to lack nitrile-specifier activity (Kong et al., 2012), it has a partially conserved putative iron-binding triad (Brandt et al., 2014). As a result, it has been proposed that *At3g07720* is the ancestor to *NSPs* and may function in iron uptake, transport, or storage (Brandt et al., 2014). We therefore tested whether insertions in *ISR1* resulted in a growth defect when plants were grown under iron limitation, but we found no difference in root elongation under low iron conditions relative to the wildtype controls (Supplementary Fig. 3). We also performed a ferric chelate reductase assay to measure the reduction of Fe(III) to Fe(II) by the root FRO2 enzyme during the iron deficiency response (Yi and Guerinot, 1996). The *frd3* loss of function mutant, which exhibits constitutive ferric chelate reductase activity was used as a control (Rogers and Guerinot, 2002). We found that the loss of *ISR1* function had no effect on ferric chelate reductase activity (Supplementary Fig. 3). These data indicate that although *ISR1* is induced under iron deficiency conditions, the loss of *ISR1* is not essential for plants to obtain and utilize iron during the iron deficiency response.

To test if *ISR1* contributes to plant iron levels or iron localization, we quantified total and apoplastic iron. We did not find a significant change in shoot and root levels of iron in the Col-0 *isr1-1* mutant by ICP-MS analysis (Fig. 2A-B). We used *frd3*, which has significantly higher levels of iron in shoots as a control (Rogers and Guerinot, 2002). Next, we looked at subcellular localization of iron to determine if *isr1-1* affects iron localization. Indeed, we found a significant increase in the apoplastic iron levels in the *isr1-1* mutant (Fig. 2C). This suggests that while the total iron levels in *the isr1-1* mutant are not altered, *ISR1* plays a role in iron distribution.

**Figure 2.**
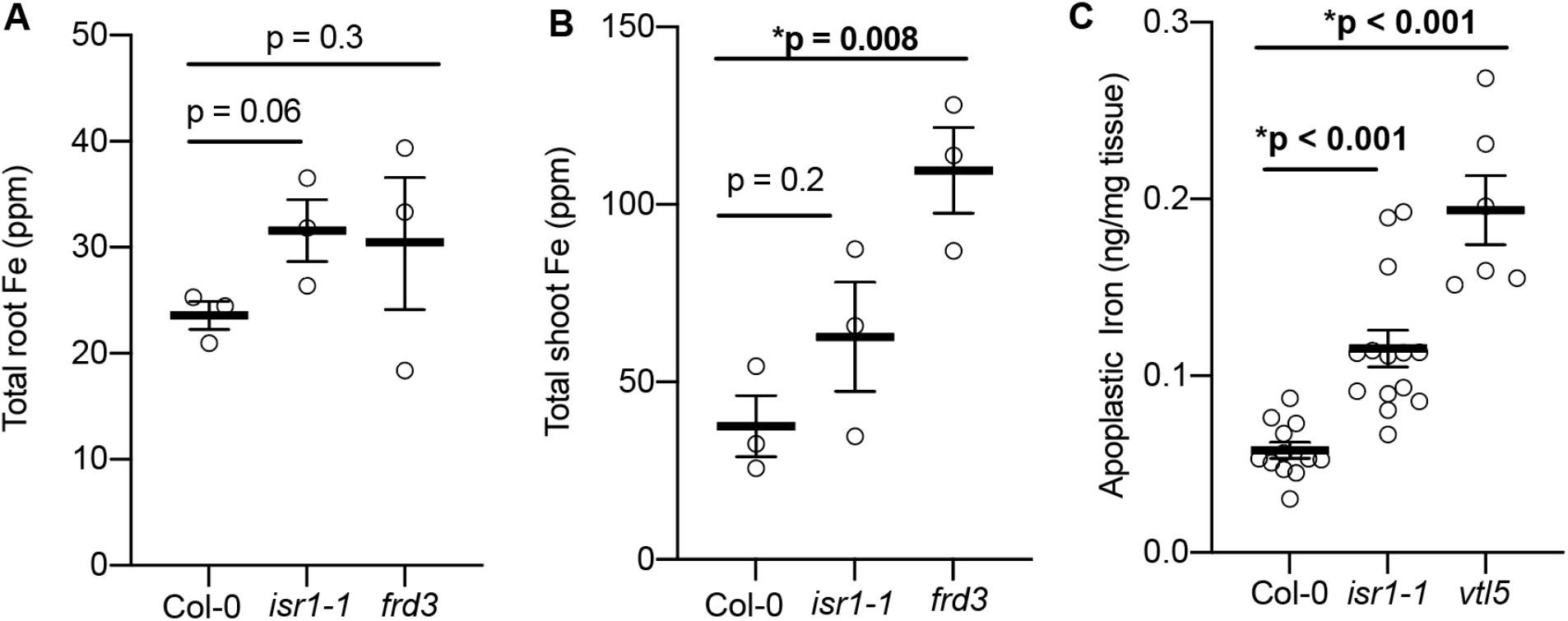
Loss of *ISR1* results in increased levels of apoplastic iron in the root. A-B, The *isr1-1* mutant did not differ from Col-0 in (A) shoot or (B) root levels of iron as determined by ICP-MS. C, the *isr1-1* mutant has increased levels of apoplastic iron. p-values by t-test relative to Col-0 are shown.

### Iron sufficiency promotes immunity via *ISR1*

The induction of an iron deficiency response in roots triggered by ISR bacteria promotes plant iron uptake (Zamioudis et al., 2015). To directly test the role of iron availability in plant disease resistance, we watered plants with 100 µM Fe (III)-EDTA or 300 µM of the Fe-chelator 2,2’-bipyridyl at 24 hours and 1 week before foliar infiltration with *Pto* DC3000. We found that soil supplementation with Fe (III)-EDTA (or inoculation with *P. simiae* WCS417 bacteria as a positive control) significantly decreased pathogen growth in Col-0 leaves (Fig. 3A). Conversely, depleting bioavailable iron by watering soil with 2,2’-bipyridyl augmented foliar bacterial proliferation compared to control plants (Fig. 3A). We next tested whether the decreased growth of *Pto* DC3000 caused by amending the soil with iron was dependent on *ISR1*. We found that treatment of the *isr1-1* mutant with Fe (III)-EDTA or WCS417 did not enhance resistance to *Pto* DC3000 (Fig. 3A) and that soil application of 2,2’-bipyridyl increased susceptibility of *isr1-1* (Fig. 3A). This indicates that soil iron availability positively correlated with plant resistance to *Pto* DC3000 in an *ISR1-*dependent manner.

**Figure 3.**
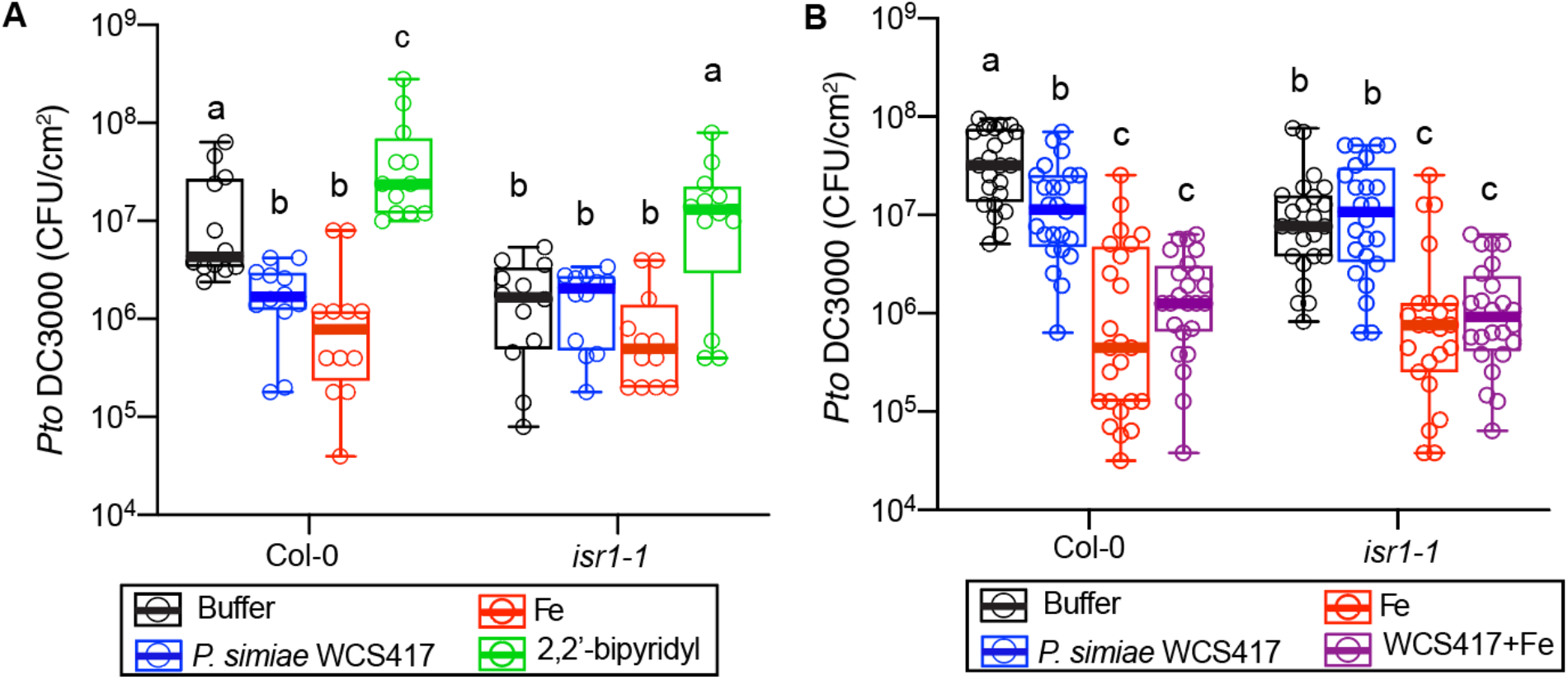
Increasing soil or apoplastic iron levels promotes immunity via *ISR1*. A, 5-week-old plants were watered with water or water supplemented with 100 µM Fe (III)-EDTA, or 300 µM of the iron chelator 2,2’-bipyridyl (2,2’-BP) 1 week and 24 hours prior to infection with *Pto* DC3000. *P. simiae* WCS417 was added to the soil 4 weeks prior to infection. CFUs/cm^2^ of *Pto* DC3000 in leaf tissue was measured 2 days after inoculation. B, Iron infiltration in leaves enhances immunity non-additively with root inoculation with WCS417. Plants were grown as in A, but 24-hours prior to infection, leaves were infiltrated with 50 µM Fe (II)SO_4_ or water. A-B, Letters indicate statistical significance p<0.01 by ANOVA and Tukey’s HSD tests (n=12 leaves for each treatment from two separate experiments)

Since we found an increase in apoplastic iron in an *isr1* loss-of-function mutant, we hypothesized that ISR bacteria might increase the availability of iron in shoots in an *ISR1*-dependent manner. To test whether iron and *P. simiae* WCS417 act through *ISR1* to enhance immunity, we infiltrated 50 µM Fe (II)SO_4_ or water into the leaves of wild type Col-0 or *isr1-1* soil-grown plants that had been grown in the presence of *P. simiae* WCS417 or buffer. We used Fe (II) for these experiments since it can directly participate in the Fenton reaction. One day after leaf infiltration with 50 µM Fe (II)SO_4_, we challenged leaves with *Pto* DC3000. We found that iron infiltration into leaves of wild-type Col-0 plants enhanced immunity nearly 1000-fold against *Pto* DC3000, and that iron infiltration in leaves combined with root WCS417 treatment resulted in similar levels of resistance as iron infiltration alone (Fig. 3B). Additionally, we found that iron enhanced resistance in the *isr1-1* mutant, but to a lesser extent (∼100-fold in *isr1-1* versus ∼1000-fold in Col-0) than the enhanced resistance observed in Col-0 (Fig. 3B). Additionally, we saw no additive effect by *P. simiae* WCS417 and iron in the *isr1-1* mutant, similar to wild-type plants. Collectively, these data indicate that WCS417 cannot further enhance immunity in the presence of high apoplastic iron suggesting that WCS417 and iron promote immunity through the same mechanism. Additionally, as we found that exogenous iron does not promote resistance in an *isr1* mutant to the same degree as in wildtype plants, indicating that iron promotes immunity in a partially *ISR1*-dependent manner.

### *ISR1*, iron, and *P. simiae* WCS417 non-additively enhance the defense-triggered ROS burst

We hypothesized that the increase in apoplastic iron in the *isr1* mutant may directly contribute to immunity through the Fenton reaction resulting in higher ROS in the apoplast. To test this hypothesis, we infiltrated leaves of Col-0, *isr1-1* or the *isr1-1; 35S:ISR1-GFP* genotypes with 50 µM Fe (II)SO_4_ 24 hours prior to performing ROS burst assays. Plants were grown with or without rhizosphere treatments of *P. simiae* WCS417 and leaf disks were used for ROS burst assays. Following exposure to the synthetic peptide flg22 corresponding to bacterial flagellin that elicits a strong ROS response (Felix et al., 1999; Gómez-Gómez et al., 1999), we found that Col-0 leaves infiltrated with iron had enhanced apoplastic ROS production indicating that increased apoplastic iron can directly enhance the ROS burst (Fig. 4A). Root treatment with *P. simiae* WCS417 also resulted in enhanced ROS (Fig. 4A). Interestingly, we saw no additive effect of *P. simiae* WCS417 and iron in the magnitude of the burst (Fig. 4A), which is consistent with the hypothesis that WCS417 may contribute to defense by promoting iron uptake.

**Figure 4.**
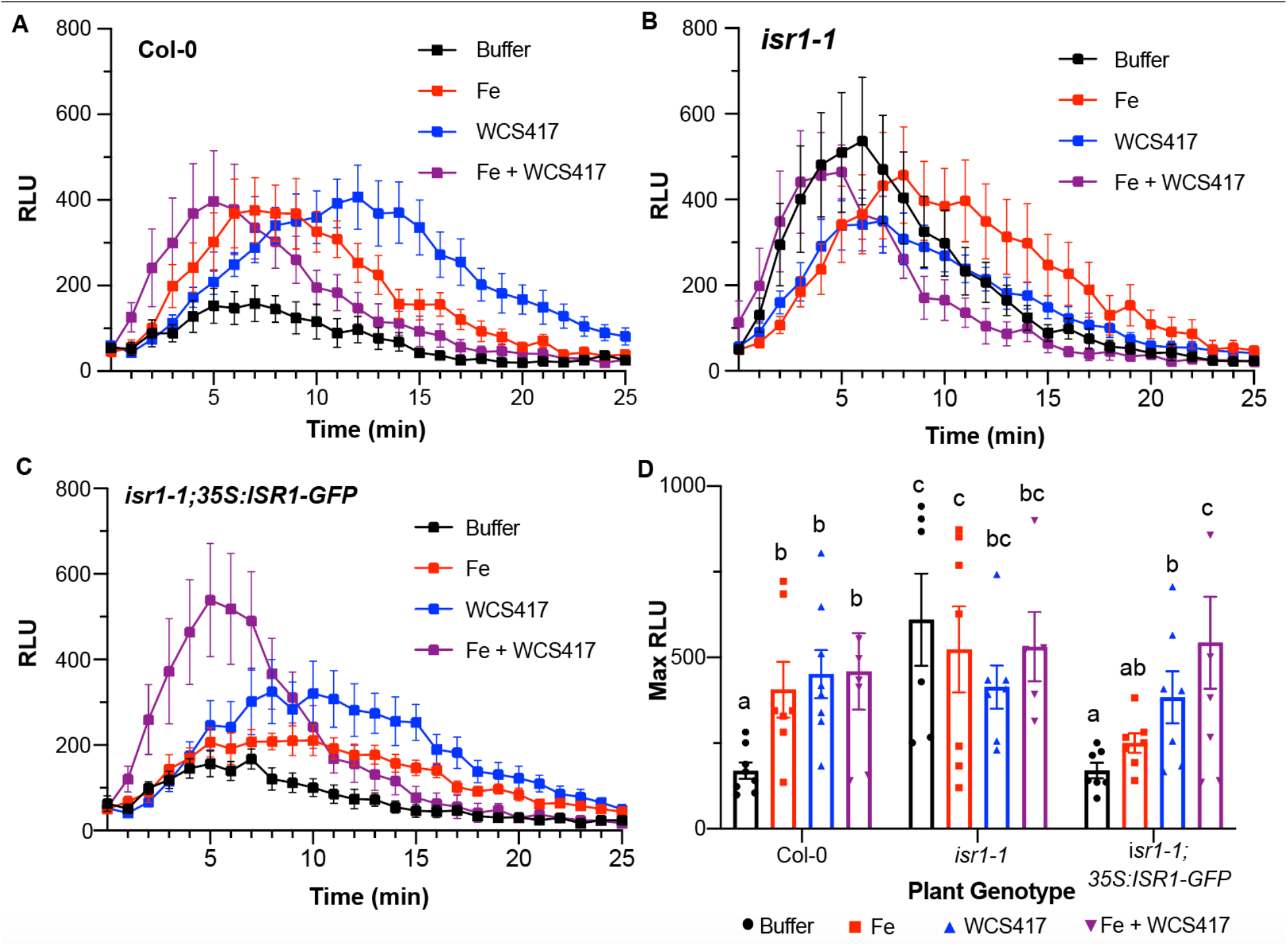
Iron, ISR bacteria and *ISR1* enhances apoplastic ROS formation. A-C, Leaf disks were extracted from 4-week-old soil-grown plants and used in ROS-burst assays. 24 hours prior to leaf disk harvest, leaves were infiltrated with water or 100 µM Fe (II)SO_4_. Values represent mean Relative Light Units (RLUs) over time. n=8 leaf disks/treatment. The experiment was repeated twice with similar results. The assay was performed with A, Col-0, B, *isr1-1* and C. The *35S:ISR1-GFP* overexpression line. D, Select time point (8 min) comparing ROS production in WT, *isr1-1*, and *35S:ISR1-GFP* seedlings. Letters designative p<0.05 by ANOVA and Tukey’s HSD test; SEM is shown.

To determine if *ISR1* is necessary for iron and *P. simiae* WCS417-mediated enhancement of leaf ROS, we repeated the above experiment with the *isr1-1* and *isr1-1; 35S:ISR1-GFP* lines. We found that *isr1-1* leaf punches had a significantly enhanced ROS burst compared to WT Col-0 leaf tissue (Fig. 4B). However, in the *isr1-1* mutant, neither iron, WCS417, or combined iron and WCS417 treatments resulted in further enhancement of ROS production. In contrast, the *isr1-1; 35S:ISR1-GFP* line had a similar level of ROS production as Col-0 that could be enhanced by WCS417 or iron treatment (Fig. 4C). Collectively, these data suggest that iron, *P. simiae* WCS417 and *ISR1* regulate plant defense-induced ROS by modulating apoplastic iron levels.

## Discussion

We found that the iron-regulated gene, *At3g07720*, encodes the *ISR1* gene (Fig. 1C). Loss of function *isr1* mutants were significantly more resistant to infection by *Pto* DC3000 whereas overexpression lines were highly susceptible (Fig. 1C), indicating ISR1 plays a role in immunity. We discovered that plant resistance to *Pto* DC3000 correlated with enhanced ROS formation in the *isr1-1* mutant (Fig. 4). ROS formation is one of the first signs of the activation of a plant innate immune response, and successful infection of a pathogen is typically dependent on an initial weak defense response of the plant to the invader (Qi et al., 2017). Because the oxidative burst is one of the first lines of defense against many biotic stresses, apoplastic iron levels may play a role in determining ROS production during defense against *Pto* DC3000. The *isr1-1* mutant generates more ROS than WT, presumably due to the increased apoplastic iron contributing to ROS production.

Iron plays a dual role in plant immunity: low iron availability can restrict pathogen growth whereas excess iron can contribute to generation of reactive oxygen species (Liu et al., 2007; Segond et al., 2009; Aznar et al., 2014; Ye et al., 2014; Aznar and Dellagi, 2015; Xing et al., 2021). Because *isr1* mutants do not display a phenotype on iron-deficient medium, it is unlikely that *ISR1* is essential for plant viability under iron limitation. Due to the potential toxicity of iron to the cell, it is essential that plants are able to acquire adequate nutrition without promoting tissue damage. Therefore, we speculate that *ISR1* is upregulated during the iron deficiency response to protect plant cells against ROS during the sudden influx of iron into the apoplast. *ISR1* has similarity to *Arabidopsis* nitrile-specifier proteins (NSPs), which use iron as a cofactor (Kuchernig et al., 2012). Although *ISR1* lacks nitrile-specifier activity (Kong et al., 2012), it has a partially conserved putative iron-binding triad (Brandt et al., 2014). ISR1 may function to bind iron inside the cell, or another subcellular compartment, sequestering it away from the apoplast.

Our data suggest that iron may contribute to the systemic signal to prime plants for pathogen attack during ISR. The upregulation of iron-uptake genes by WCS417 bacteria can rescue chlorosis and other signs of iron starvation through enhanced uptake (Zamioudis et al., 2015; Trapet et al., 2016). We hypothesized that WCS417 bacteria activate the iron deficiency response to accumulate iron for enhanced defense. *ISR1* is a negative regulator of this process; it may be induced during times of iron uptake to avoid toxicity from excess apoplastic iron. Future work on the function of *ISR1* will elucidate how iron homeostasis is coordinated with defense signaling to regulate plant immunity.

## Materials and Methods

### Identification of candidate genes within the ISR1 locus

The *ISR1* locus was previously mapped to chromosome 3 between the B4 (2.25 Mb) and GL1 markers (10.36 Mb) (Ton et al., 1999). Candidate genes were identified as those between markers B4 and GL1 that had significant induction of at least log_2_ = 1.5 in response to ISR or iron deficiency in published datasets (Dinneny et al., 2008; Zamioudis et al., 2015) (Supplementary Table 1).

### Ws genome sequencing

DNA from Ws seedlings was extracted and sent for PE150 Illumina sequencing (Novogene). Reads were mapped to the Col-0 genome using the Integrative Genomics Viewer (Robinson et al., 2011) and the 8 candidate genes were individually queried for non-synonymous mutations. The genomic rearrangement in *At3g07720* was validated by PCR using primers shown in Supplementary Table 2.

### Plant Materials and Growth Conditions

The *Arabidopsis* T-DNA insertion lines corresponding to *ISR1* in the ecotypes Col-0 and Ws, *isr1-1* (SALK_012767.35.00X) and *isr1-2* (Flag_436E03) respectively, were obtained from the *Arabidopsis* Biological Resource Center (ABRC) (https://abrc.osu.edu). The *myb72* mutant is a T-DNA line (SALK_052993) that was previously characterized (Palmer et al., 2013). Homozygous lines were identified by PCR using gene-specific (*ISR1* start FOR and *ISR1* stop REV) and T-DNA-specific left border primers (Salk LBb1.3 FOR and *ISR1* stop REV for *isr1-1* lines; Tag5 FOR and *ISR1* stop REV for *isr1-2* lines; Table S1). For soil-grown plants, seeds were surface sterilized in 70% ethanol and a solution of 25% bleach and 0.2% SDS. Seeds were then stratified and imbibed in sterile distilled water for 3 days and sown directly onto Jiffy-7 peat pellets (Jiffy products) and grown at 12-hour light, 12-hour dark, 22**°**C with low light (75-100 µmol m^-2^ sec^-1^). For plate grown plants, seeds were surface sterilized using vapor-phase sterilization for 4-12 hours and plated on agar medium and stratified/imbibed for 3 days. Plants were grown on 0.5X Murashige and Skoog (MS) medium (Sigma) with 1% sucrose, 1mM MES and 1% Agar Type M (Sigma), pH adjusted to 5.7 with KOH. Plants were grown at 16-hour light, 8-hour dark, 22**°**C.

### Plasmid construction and plant transformation

#### pCR8 entry vector constructs

*ISR1* CDS was amplified using Go-Taq (Promega) with *ISR1* start FOR and *ISR1* no stop REV primers (Table S1). *ISR1* CDS with a 5’ EcoRI restriction site and 3’ BamHI restriction site was amplified using Phusion (Thermo Fisher Scientific) and *ISR1* start + EcoRI FOR and *ISR1* no stop + BamHI REV primers (Table S1). Template cDNA was generated from Col-0 plants grown on iron-free media. CDS was subcloned into pCR8 using pCR8/GW/TOPO TA cloning kit (Thermo Fisher Scientific). All final constructs were sequenced using vector and gene specific primers to ensure no mutations and correct alignment (See Table S1 for all sequencing primers).

#### 35S overexpression construct

The pCR8-*ISR1* construct was used to subclone the gene into the pMDC32 (35S cauliflower mosaic virus [CaMV] promoter) or pEarleyGate103 (35S CaMV promoter with an N terminal GFP fusion) vectors using Gateway LR Clonase II (Thermo Fisher Scientific) and then stably expressed into *isr1-1* and *isr1-2* plants following *Agrobacterium tumefaciens* (strain GV301)-mediated transformation using the floral dip method (Clough and Bent, 1998).

### ISR infection assays

ISR assays were performed as described (Cecchini et al., 2019). WCS417 was cultured overnight at 28°C in King’s B medium containing 50 µg/ml of rifampicin. Cells were diluted to a final OD_600_ of 0.05 (5×10^5^ CFU/g soil) in 10 mM MgSO_4_ buffer. Each Jiffy pellet was inoculated with 2 ml of diluted WCS417 bacterial strain 9 days after seed germination. Control plants were inoculated with 2 ml of 10 mM MgSO_4._

### *P. syringae* pv *tomato* DC3000 infection assays

*Pto* DC3000 was cultured overnight at 28°C in King’s B medium containing 50 µg/ml of rifampicin. Bacterial suspension was diluted to OD_600_ of 0.01 then to a final OD_600_ of 0.0002 (2×10^3^ CFU/ml) in 10 mM MgSO_4._ Plants were grown in Jiffy-7 peat pellets under a 12 h light/12 h dark and a 23/20 °C day/night temperature regime and 75-100 µmol m^-2^sec^-1^ light. *Pto* DC3000 infection assays were performed and analyzed as described (Cecchini et al., 2019). Infections were performed on mature leaves of 5-week-old plants.

For iron treatment, 1 ml of 0.1 mM Fe (III)-EDTA was pipetted directly onto Jiffy pellets 1 and 7 days prior to *Pto* DC3000 infection. Distilled water was used a control.

### Quantitative Real Time PCR

Plants were grown on 0.5X MS media for 10 days and transferred to 50 µM iron or 300 µM ferrozine plates for 3 days. Root and shoot tissue were separated and snap frozen in liquid nitrogen. Total RNA was isolated using the RNeasy kit (Qiagen) and treated with DNaseI (Roche). 1 µg of RNA was used for reverse transcription by the M-MLV-RT (Life Technology) using oligo(dT) primer and qRT-PCR was performed on a Step One Plus system (Applied Biosystems Version 2.2.3) using SYBR Premix ExTaq reagents and protocol (Takara). Experiments are performed with 3 technical and biological replicates. Relative transcript levels were calculated by normalizing to *clatherin* expression. The primers used are listed in Table S1.

### Ferric Chelate Reductase Assay

Ferric Chelate Reductase assays were performed as described (Yi and Guerinot, 1996). Plants were grown on 0.5X MS media for 10 days and transferred to 50 µM iron or 300 µM ferrozine plates for 3 days before roots were assayed.

### Apoplast iron measurement assays

Apoplast iron measurement assays were performed as described (Frits Bienfait et al., 1985). Plants were grown on 1X B5 media with 100 µM iron for 10 days before roots were tested.

### Ethylene sensitivity assays

Ethylene sensitivity assays were performed as described (Ton et al., 2001). For ACC treatment, plants were imbibed and stratified at 4°C for 3 days on 0.5X MS media -iron with 0.5% sucrose. ACC was added to a final concentration of 1µM from a 1mM stock of ACC resuspended in water with 0.96% (v/v) ethanol and filter sterilized with a 0.22 filter. Control plates were absent of ACC.

### Apoplast chemiluminescence assays

For leaf disc assays, plants were grown for 5 weeks at 12-hour light/12-hour dark, 22**°**C in Jiffy pellets. For iron treatment experiments, 24 hours prior to leaf extraction 50 µM Fe (II)SO_4_ or water was infiltrated into leaves. A 0.5 cm diameter cork borer was used to remove equal sized discs from mature leaves. Leaf discs were cut into 4 equal sized pieces to increase exposed apoplast surface area. Half of each leaf disc was placed abaxial side up in a single well of a white 96-well plate containing 100 µl water and allowed to recover overnight in plant growth chamber. One hour before elicitation plates were placed in the dark. Flg22 (PhytoTechnology Laboratories) was prepared fresh from stock solutions stored at −20°C and used as the elicitor for all experiments. Flg22 was added at a final concentration of 100 nM. The luminol assay was performed in the dark as described by (Mammarella et al., 2015). A M1000 PRO Tecan plate reader was used to record relative light units (RLUs) every two minute for 30 minutes.

### Elemental Analysis

For elemental analysis of soil grown plants, seeds were sown in Metro Mix 820 and grown for 3 weeks. Whole rosettes from 6 plants per genotype were collected and washed in 2mM CaSO_4_ followed by 5mM EDTA and distilled water to remove trace metals then dried overnight at 60°C. Plate grown plants were grown on 10µM Fe for 10 days then transferred to either 10µM Fe or 50µM Fe supplemented with 10µM CoCl_2_. Plants were exposed to 10µM Fe instead of 5µM Fe so WT and knockout mutant plants are the same size. Shoot and root tissue were separated from roots with a razor and washed in 2mM CaSO_4_, 5mM EDTA and distilled water then dried overnight at 60°C. ICP-MS analysis was performed on these tissues at the University of Aberdeen as previously described (Baxter et al., 2007).

### Statistical analyses

ANOVA followed by Tukey HSD were performed to calculate statistically significant differences between data sets. A p value of 0.05 was considered significant

## Data availability

Reads from the Ws genome will be made available upon publication.

## Author contributions

A.L.S., F.M.A., M.L.G., and C.H.H. conceived of the study. A.L.S. characterized gene expression levels and iron quantification in the *isr1* mutants and overexpression lines. A.L.S. and B.R. quantified apoplastic iron. Y.S., J.B., and C.H.H performed ISR, immunity and ROS experiments. A.L.S., Y.S., M.L.G. and C.H.H. analyzed data. A.L.S. and C.H.H wrote the manuscript with input from all. C.H.H. and M.L.G. agree to serve as the author responsible for contact and ensure communication.

## Acknowledgements

This work was supported by an NSERC Discovery Grant (NSERC-RGPIN-2016-04121), a Seeding Food Innovation grant from George Weston Ltd., a prior Tosteson Fund for Medical Discovery postdoctoral fellowship to C.H.H, by NIH R37 grant GM48707 and NSF grant IOS-0929226 awarded to F.M.A and by NSF grants IOS-1456290 and IOS-1257722 to M.L.G. Y.S. was supported by a fellowship from the China Postdoctoral Science Foundation. We thank Alicia Sivitz for constructing the transgenic overexpression lines and David Salt and John Danku for generating the ICP-MS data.

## Supplemental Figures and Tables

**Supplemental Figure 1.**
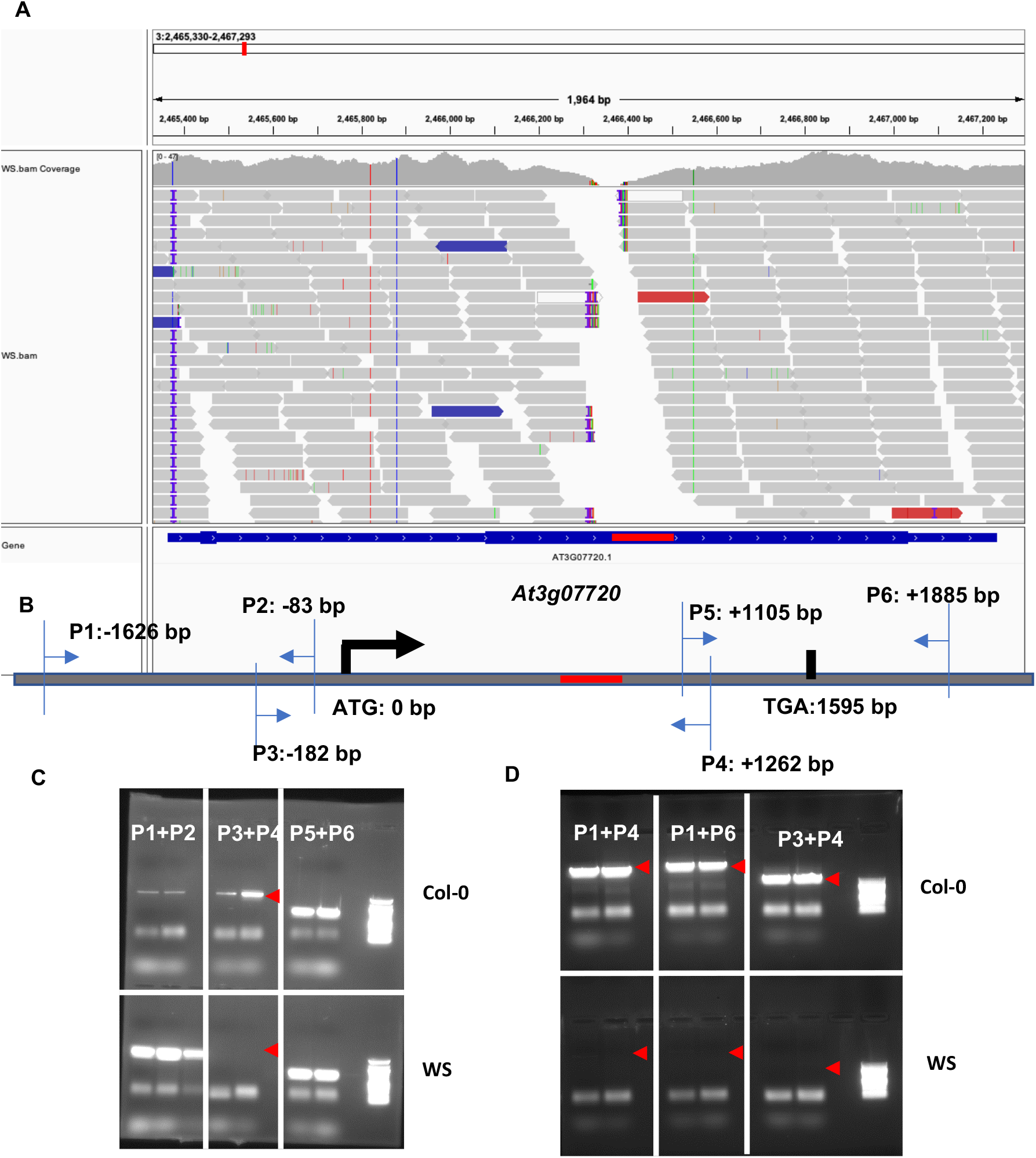
A genomic rearrangement or insertion likely occurred within the Ws *At3g07720* gene. A, Illumina sequencing of the Ws genome revealed a ∼75 bp region with no mapped reads without the *At3g07720* genome. B-D, Attempts to PCR through the region in Ws were unsuccessful suggesting that rather than a small deletion at this locus, that Ws has a large insertion or rearrangement.

**Supplemental Figure 2.**
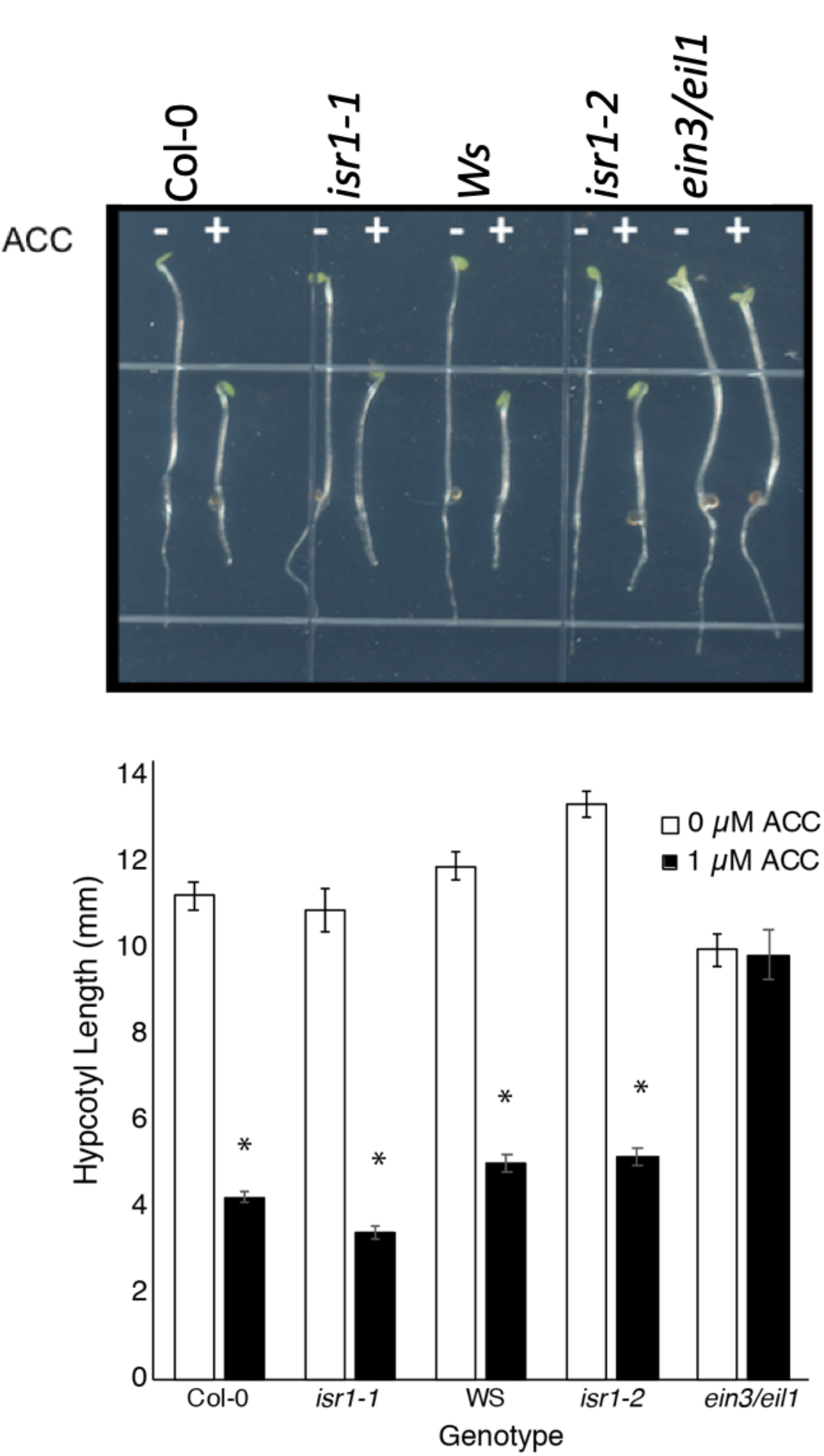
*ISR1* loss-of-function mutants are not ethylene insensitive. The triple response was assessed in Col-0, *isr1-1*, Ws and *isr1-2. Ein3/eil1* ethylene insensitive mutants were used as a positive control. Ethylene-insensitive plants will not exhibit decreased hypocotyl or root elongation or an apical hook in the presence of ACC, unlike the mutant and WT lines. Seeds were plated on 0.5X MS agar plates +/- 1µM of the ethylene precursor 1-Aminocyclopropane-1-carboxylic acid (ACC). Experiment was repeated 3 times with similar results. n>20; *<0.01 by t-test relative to untreated control of the same genotype.

**Supplemental Figure 3.**
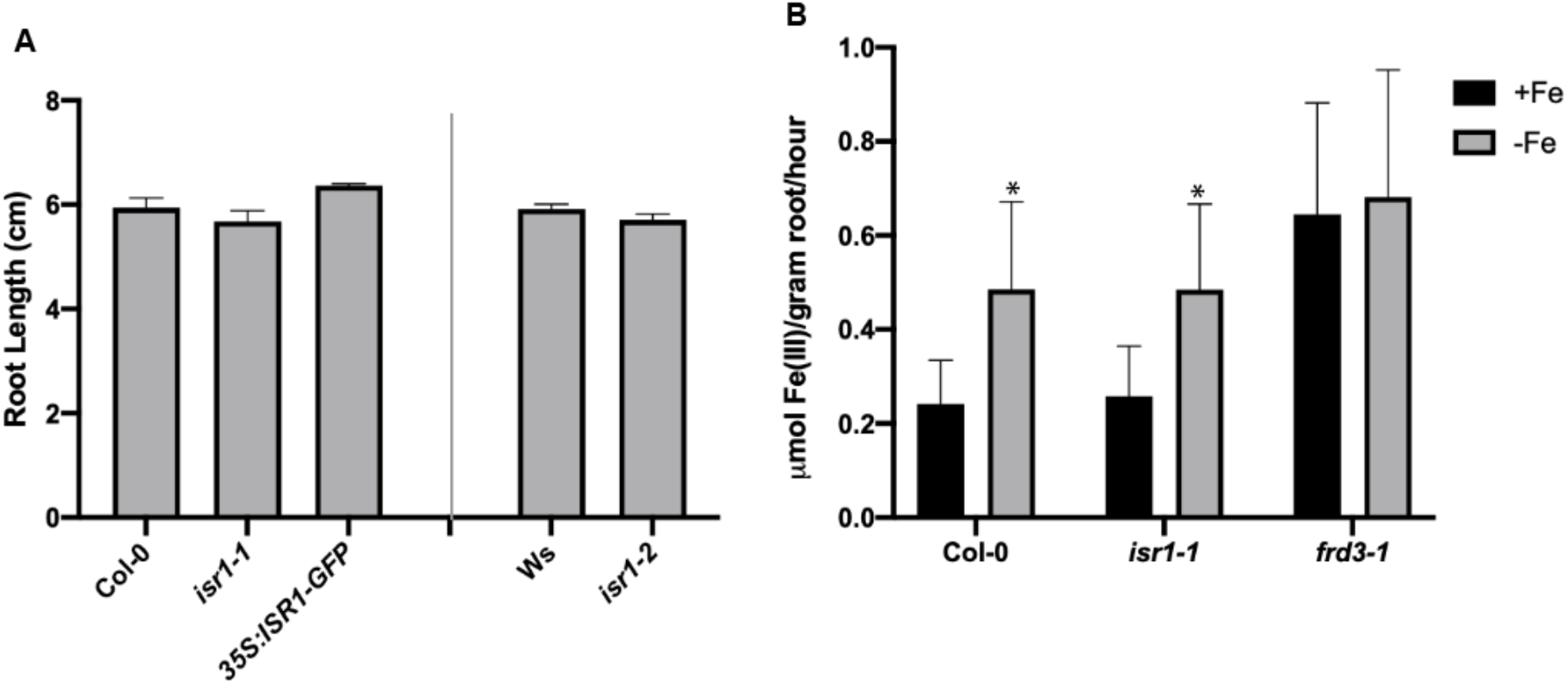
Root growth in iron deficient conditions and Ferric chelate reductase activity is unaffected by insertion in *ISR1*. A, Root length of Col-0, Ws, *At3607720-1* and Ws *At3g07720-2, At3g07720* overexpression in Col-0 were grown under iron limiting conditions for 14 days. B, Plants were grown on 0.5X MS for 10 days and transferred to media with 50 µM iron (+Fe) or 300 µM Ferozzine (-Fe) for 3 days. While Col-0 *frd3* shows constitutive activity of FRO2 reductase regardless of iron status, there was no change in activity in Col-0 or *isr1-1*. *<0.05 by t-test relative to +Fe control of the same genotype

**Supplementary Table 1.** Candidate genes within the ISR1 locus. The *ISR1* locus was previously mapped to chromosome 3 between the B4 (2.25 Mb) and GL1 markers (10.36 Mb) (Ton et al., 1999). Candidate genes were identified as those between markers B4 and GL1 that had significant induction of at least log_2_ = 2 in response to ISR or iron deficiency. Previously described genes required for ISR *MYB72* and *BGLU42* are outside of the ISR1 locus but included for reference. Whether the gene has predicted non-synonymous mutations based on sequencing Ws is indicated.

| Gene ID | Predicted Function | Log<sub>2</sub> FC (ISR) | Log<sub>2</sub> FC (Iron deficiency) | Mutations in Ws |
| --- | --- | --- | --- | --- |
| At3g07720 | Galactose oxidase/kelch repeat superfamily protein | 5.71 | 33.79 | $\Delta 93-116$ |
| At3g10720 | Plant invertase/pectin methylesterase inhibitor superfamily | 3.43 | 4.17 | M1I, H36R, F519L |
| At3g12820 | AtMYB10, MYB10, MYB domain protein 10 | 2.77 | 4.67 | N |
| At3g12900 | sh8, 2-oxoglutarate (2OG) and Fe(II)-dependent oxygenase superfamily protein | 42.32 | 26.25 | N |
| At3g13610 | f6'h1, 2-oxoglutarate (2OG) and Fe(II)-dependent oxygenase superfamily protein | 4.07 | 36.06 | N |
| At3g18250 | Putative membrane lipoprotein | 10.07 | 5.33 | N |
| At3g18560 | unknown protein | 2.12 | 4.12 | N |
| At3g26830 | CYP71B15, PAD3, Cytochrome P450 superfamily protein | 14.78 | 4.18 | N |
| At1g56160 | MYB72 (MYB DOMAIN PROTEIN 72); DNA binding / transcription factor | 10.22 | 4.54 | N |
| At5g36890 | BGLU42 (BETA GLUCOSIDASE 42); beta-glucosidase/ catalytic/ cation binding / hydrolase, hydrolyzing O-glycosyl compounds | 4.51 | 15.28 | N |

**Supplemental Table 2.**
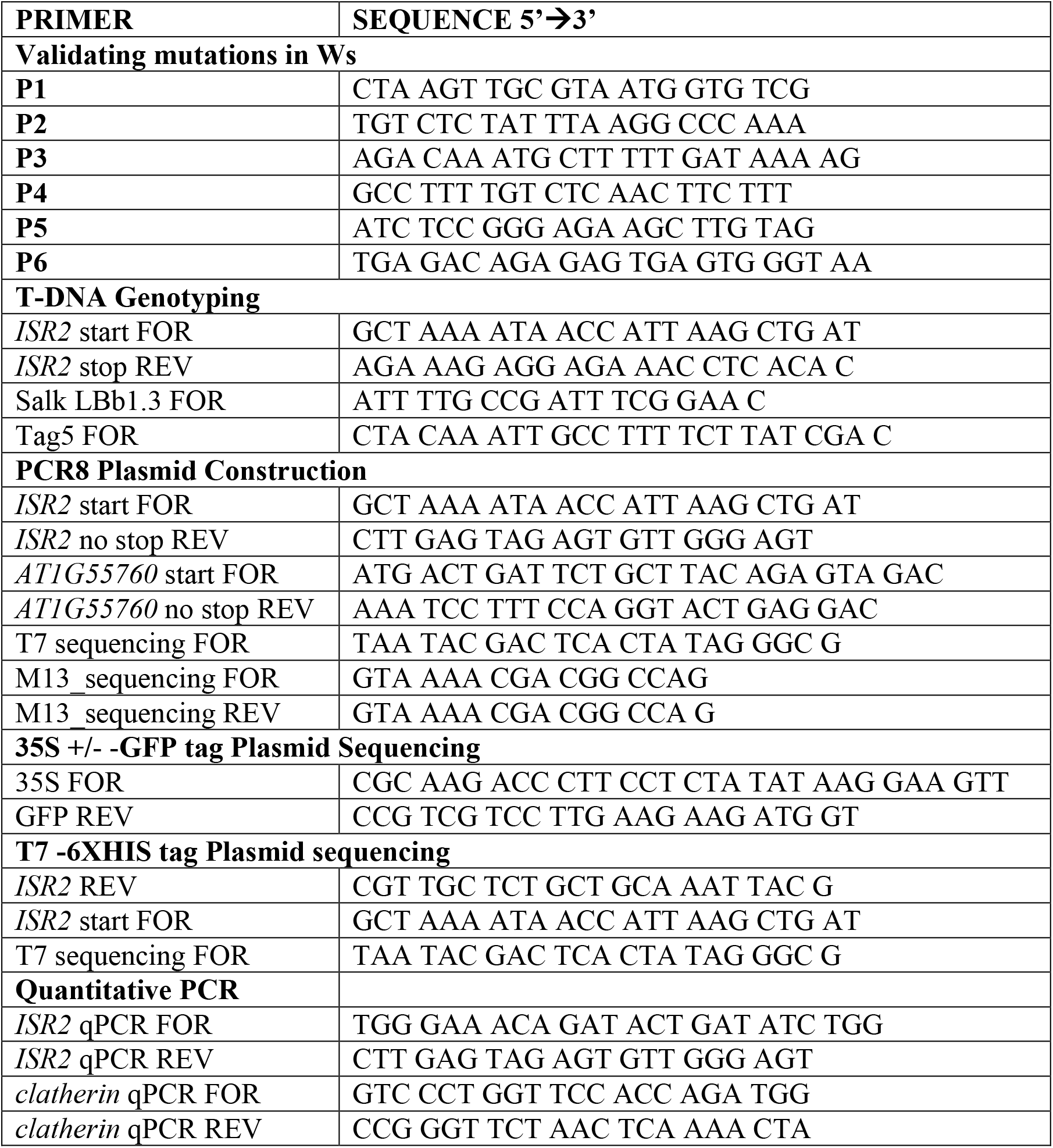
Primers used in this study.

